# Genomics of eye number evolution in spiders

**DOI:** 10.1101/2025.07.21.665793

**Authors:** Chao Tong, Zheng Fan, Luyu Wang, Zhisheng Zhang

## Abstract

Spiders exhibit tremendous variations in eye numbers, but the genomic basis underlying this diversity remains largely unexplored. Here, we analyzed the genomic data of 148 spider species in 31 families, representing all phenotypes of eye numbers including 8, 6, 4, 2, and 0. Our analyses revealed that a core set of 29 genes involved in eye development and phototransduction is conserved across all spiders, irrespective of their eye number, indicating that eye reduction is not caused by the loss of key developmental genes. We found that evolutionary transitions leading to reduced numbers of eyes are primarily associated with parallel genome-wide relaxed selection. While these independent reduction events shared several genes under consistent selective pressures, they did not share any genes under positive selection, indicating putatively divergent molecular mechanisms. In contrast, the evolution of complete eye loss in cave-dwelling spiders is associated with parallel genome-wide intensified selection. Notably, we identified shared genes under both intensified and positive selection across independent origins of eyelessness, suggesting a parallel molecular mechanism. Altogether, our study provides genomic insights into the parallel evolution of eye reduction and complete eye loss in spiders.

## Introduction

The evolution of spider visual systems is fascinating but complex, characterized by tremendous diversity in eye number (e.g. 8, 6, 4, 2, 0 eyes), eye arrangement and eye size (1–3). For example, most spider species have 8 eyes, while few species have no eyes (Fig. 1A). This raises the intriguing question: why do spiders have diverse eye numbers? Past studies have focused on understanding how environmental factors and ecological niches shape eye number evolution (2). For instance, cave-dwelling spiders exhibit reduced or no eyes, representing a trait associated with adaptation to lightless habitats (4–8). Thus, spiders in visually complex environments typically possess diverse visual systems, facilitating enhanced adaptive capacity.

**Figure 1.**
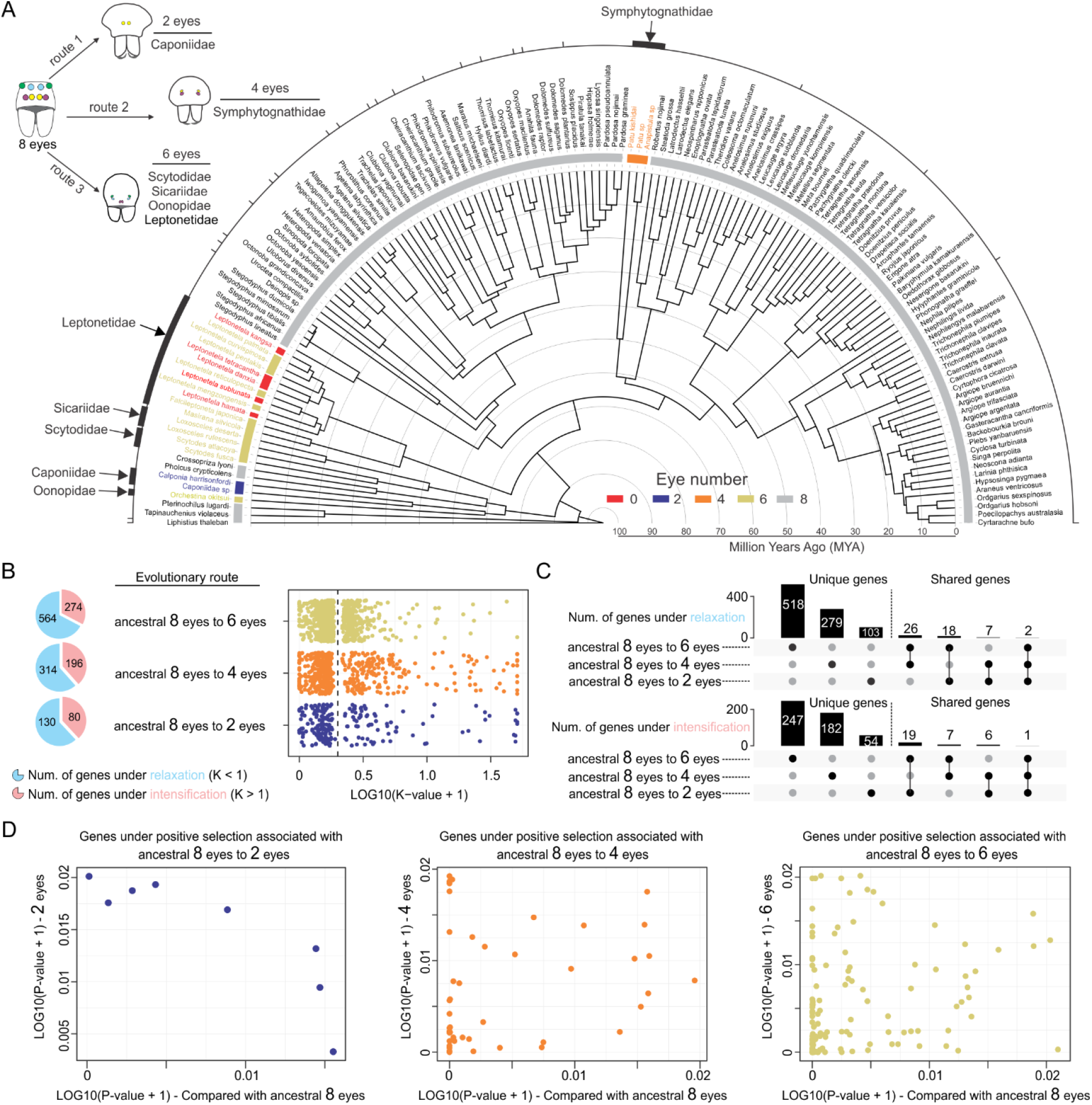
Evolutionary routes and associated genetic changes of eye reduction in spiders. (A) A time-calibrated phylogeny of 148 spider species based on 1,827 BUSCO single-copy orthologous genes. Species with 0, 2, 4, 6, and 8 eyes are indicated by red, blue, orange, yellow, and grey colors, respectively. The tree highlights three independent evolutionary routes of eye reduction from an eight-eyed ancestor: to 2 eyes (Route 1), to 4 eyes (Route 2), and to 6 eyes (Route 3). (B) Pie charts depicting the numbers of genes under relaxed (light blue) or intensified (pink) selections associated with each route. Scatter plot depicting the distribution of selection intensity (K-values from RELAX analysis) for these genes. (C) UpSet plots illustrating the number of unique and shared genes among the three routes. The top plot shows genes under relaxed selection, and the bottom plot shows genes under intensified selection. (D) Scatter plots depicting the genes under positive selection associated with each evolutionary route leading to eye reduction. Each dot represents a gene identified by a branch-site model test in BUSTED-PH. The Y-axis indicates the log-transformed P-value for positive selection on the focal foreground branches (2-eyed, 4-eyed, or 6-eyed lineages). Genes with lower P-values show stronger evidence of positive selection. The X-axis indicates the log transformed P value by comparing focal foreground branches (i.e. 2-eyed spiders) with ancestral branches of 8-eyed spiders.

Parallel evolution of eye numbers is common in spiders. One exemplar trait is the consistent eye number in phylogenetically close clades (i.e. family), such as 6-eyed species in families Scytodidae, Sicariidae, Oonopidae, and Leptonetidae (1) (Fig. 1A). Another example is the parallel complete loss of eyes in cave spiders within genus *Leptonetela* (4, 7, 8). These parallel features underscore the role of environmental pressures in shaping the evolution of eye numbers in spiders, illustrating how different species have parallelly evolved similar traits to meet the challenges of their respective ecological niches, while the underlying genomic landscape remains largely unexplored.

Here, we assembled a comprehensive genomic dataset of 148 species within 31 spider families, representing all phenotypic traits of eye numbers including 8, 6, 4, 2 and 0. Using this dataset, we aimed to uncover parallel changes associated with the evolution of eye numbers in spiders. Specifically, we first tested whether presence or absence of key genes involved in eye development and phototransduction are associated with the evolution of eye numbers in spiders. This is because a recent study identified a nearly complete set of phototransduction pathway genes in eyeless cave spiders (4), while this evolutionary scenario has not been fully tested in spiders with diverse eye numbers across the phylogeny. We further tested whether parallel changes in the patterns of molecular evolution are associated with multiple evolutionary transitions leading to the diversity of eye numbers.

## Results

We analyzed the publicly available high-quality genomes of 27 spider species with 8 eyes and representative transcriptome assemblies of 22 species with 6, 4, 2, or 0 eyes (Dataset S1). Our results showed that a core set of 29 genes associated with eye development and phototransduction is retained across all these species, irrespective of their eye numbers (Dataset S2). This comparison across species with varying degrees of eye reduction reveals that the fundamental genetic toolkit for vision and eye formation has not been lost, even in lineages where the number of eyes is significantly reduced. The consistent presence of these 29 core genes across all analyzed genomes and transcriptomes suggests their essential role is maintained throughout the spider phylogeny.

We next focused on the patterns of molecular evolution associated with evolutionary transitions leading to the diversity of eye numbers, including shifts from 8 eyes reduced to 2 eyes (i.e., Caponiidae), 8 eyes reduced to 4 eyes (i.e., Symphytognathidae), and 8 eye reduced to 6 eyes (i.e., Scytodidae, Sicariidae, Oonopidae, Leptonetidae). In these families, the number of eyes is a consistent trait, and the six-eyed spiders, in particular, are closely related phylogenetically (Fig. 1A). Given that the ancestral state for spiders is with 8 eyes, these lineages represent distinct evolutionary routes of eye degeneration in adaptation to different environments. Through comparative genomics analysis of 9,146 orthologous genes among 148 spider species, we investigated natural selection pressures associated with different routes. We found that a majority of genes experienced parallel relaxed selection across all three major evolutionary transitions (Fig. 1B, Dataset S3). Furthermore, we identified a set of genes, including Hermansky-Pudlak syndrome 1 protein, AP-3 complex subunit delta-1, exocyst complex component 4, that were subject to shared relaxed selection across different routes (Fig. 1C, Dataset S4). Interestingly, however, the genes under positive selection were unique to each independent evolutionary route, with no overlap observed (Fig. 1D, Dataset S5), potentially indicating that different molecular mechanisms underlie parallel instances of eye reduction.

Finally, we investigated the evolution of complete eye loss (i.e. eyeless) in spiders. Unlike the clade-specific patterns observed in spiders with 2, 4, or 6 eyes, the eyeless trait often differentiates at within-family (Fig. 2A and 2B) or within-genus level (Fig. 2C). For example, species within the cave-dwelling genus *Leptonetela* (family: Leptonetidae) have lost their eyes, while other species in the same genus retain 6 eyes (4, 8). A similar pattern has been seen in family Agelenidae (typically 8 eyes) spiders, where the cave-dwelling species *Troglocoelotes proximus* is eyeless (9). We conducted phylotranscriptomic and molecular evolution analyses at genus-level (*Leptonetela*) and family-level (Agelenidae) to examine these independent instances of complete eye loss. Our analysis revealed that the degeneration of eyes into a complete eyeless state was primarily associated with intensified selection on a majority of genes (Fig. 2D, Dataset S6). Notably, two independent evolutionary paths to eyelessness shared several genes under intensified or relaxed selection, such as fibroblast growth factor receptor 1 (FGF1), Krueppel-like factor 13 (Fig. 2E, Dataset S7). Further positive selection analysis also identified a shared set of positively selected genes, such as FGF1, titin, zig3 (Fig. 2F and 2G, Dataset S8), suggesting parallel molecular evolution associated with the complete loss of eyes in these distinct lineages.

**Figure 2.**
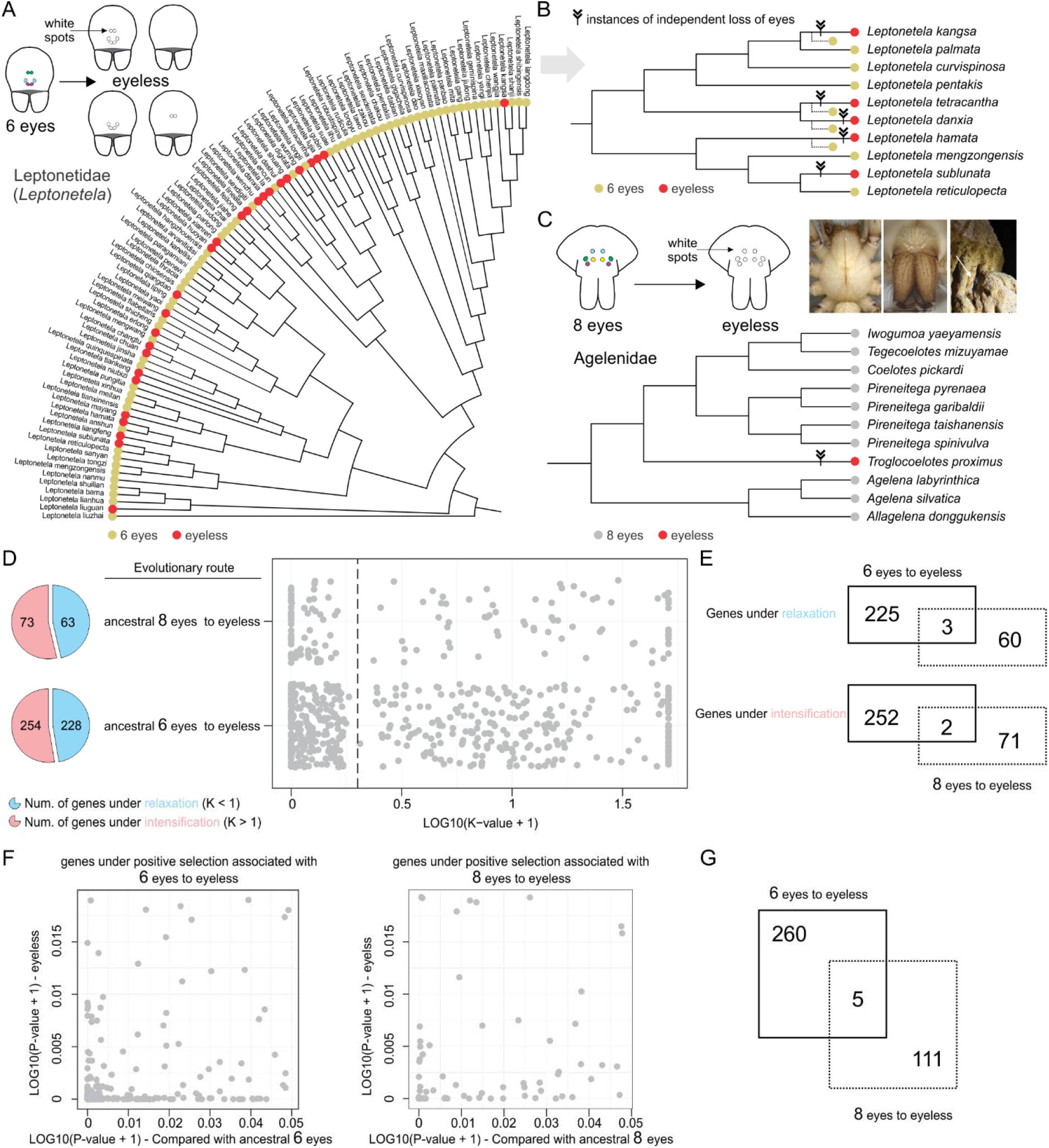
Evolutionary routes and associated genetic changes of complete eye loss in spiders. (A) Phylogeny of spiders in genera Leptonetela based on COX1 gene. Red dot indicating the eyeless species, yellow dot indicating the 6-eyed species. (B) Phylogeny of 18 Leptonetela species based on transcriptomic data. The fletch representing five instances of independent loss of eyes. (C) Phylogeny of 11 species within family Agelenidae based on transcriptomic data. Red dot indicating the eyeless species, grey dot indicating the 8-eyed species. The fletch representing unique instance of independent loss of eyes. (D) Pie chart and scatter plot depicting the genes under relaxed or intensified selection associated with parallel evolutionary routes. (E) Venn diagrams depicting the genes under relaxation (top) and intensification (bottom) shared by two parallel routes. (F) Scatter plots depicting the genes under positive selection associated with each evolutionary route leading to complete eye loss. (G) Venn diagram depicting the genes under consistent positive selection shared by two parallel routes.

## Discussion

Why do spiders exhibit such a remarkable diversity in eye numbers? One possibility is that this diversity reflects adaptations to various ecological niches they inhabit. The ancestral state for spiders is believed to be 8 eyes, and as they colonized different habitats, some lineages encountered environments with low-light conditions, such as dense bushes or leaf litter (10). In these settings, the selective pressure to maintain a full complement of eyes may have diminished, leading to various degrees of eye degeneration over extended evolutionary time. This hypothesis is consistent with our findings from the evolutionary path from 8 to 6 eyes, where we found that an eye development gene (peroxidasin) and a retinal degenerative disease gene (retinitis pigmentosa 9) experienced positive selection (11). Concurrently, we found that a gene associated with muscle development (Tropomyosin) (12) was under positive selection in these six-eyed spiders, which may be linked to compensatory adaptations in predatory behavior required in low-light environments.

Our comparative genomic analysis suggests a fundamental distinction between the evolutionary mechanisms associated with eye reduction versus complete eye loss. The transition to a reduced number of eyes appears to be a relatively moderate evolutionary process, primarily characterized by parallel relaxed selection at the genome-wide level. This “relaxed” selective pressure has allowed for diversified evolutionary outcomes, as evidenced by the fact that the positively selected genes we identified were unique and independent to each eye-reduction route. In contrast, the adaptation to the absolute darkness of cave environments imposes much stronger selective pressures. The complete loss of eyes in cave-dwelling spiders is associated with genome-wide intensified selection. This intense pressure appears to drive a more constrained and parallel evolutionary trajectory, a conclusion supported by our findings of shared intensification patterns and, critically, a common set of genes under positive selection across independent lineages of eyeless spiders.

## Materials and Methods

### Study design

Spiders exhibit tremendous diversity in number of eyes, varying from most species with 8 eyes to few species with no eyes. We sought to identify genetic basis underlying the evolution of eye numbers in spiders. We aimed to analyze and compare the genomic data of spider species representing all phenotypic traits of eye numbers including 8, 6, 4, 2, and 0.

### Omics data curation

We retrieved the genomes of spider species from online omics databases, including NCBI Genome (https://www.ncbi.nlm.nih.gov/genome), GigaDB (http://gigadb.org) and CNGBdb (https://db.cngb.org) as of May 2024. Specifically, we included the whole genomic data of *Amaurobius ferox* (GCA_951213105.1), *Argiope bruennichi* (GCF_947563725.1), *Araneus ventricosus* (GCA_013235015.1), *Argiope argentata* (GCA_026289955.1), *Argiope aurantia* (GCA_026543865.1), *Argiope trifasciata* (GCA_026543055.1), *Caerostris darwinim* (GCA_021605075.1), *Caerostris extrusa* (GCA_021605095.1), *Stegodyphus dumicola* (GCF_010614865.2), *Stegodyphus mimosarum* (GCA_000611955.2), *Hylyphantes graminicola* (GCA_023701765.1), *Oedothorax gibbosus* (GCA_019343175.1), *Pardosa pseudoannulata* (GCA_032207245.1), *Nephila pilipes* (GCA_019974015.1), *Trichonephila clavipes* (GCA_019973935.1), *Trichonephila clavata* (GCA_019973975.1), *Trichonephila inaurata* (GCA_019973955.1), *Dolomedes plantarius* (GCA_907164885.2), *Meta bourneti* (GCA_933210815.1), *Tetragnatha montana* (GCA_963680715.1), *Tetragnatha versicolor* (GCA_024610705.1), *Tetragnatha kauaiensis* (GCA_947070885.1), *Metellina segmentata* (GCA_947359465.1), *Parasteatoda tepidariorum* (GCF_000365465.3), *Latrodectus elegans* (GCA_030067965.1), *Parasteatoda lunata* (GCA_949128135.1), *Uloborus diversus* (GCF_026930045.1) (Dataset S1). Besides publicly available genomes of spiders, we also included complemental reference transcriptomes of spider species from 31 families. We employed the BUSCO pipeline (13) to assess the completeness of genome contents of these spider species based on the arachnida_odb10 single-copy orthologous gene set from OrthoDB v12 (https://www.orthodb.org), which comprised 2,934 conserved genes.

### Sample collection and RNA extraction

We collected a wolf spider species *Lycosa singoriensis* (family: Lycosidae) from Wujiaqu, China, a giant crab spider species *Heteropoda venatoria* (family: Sparassidae) from Shenzhen, China, and a jumping spider *Hyllus diardi* (family: Salticidae) from Xishuangbanna, China. In addition, we collected an eyeless species *Troglocoelotes proximus* (family: Agelenidae) from Guizhou, China. We stored all the specimens in liquid nitrogen at -80 °C, finally transferred to Southwest University, Chongqing, China. The total RNAs of whole body for each species were extracted from a single individual using Trizol (Invitrogen, USA) and glycogen (Thermo Fisher Scientific, USA) following the manufacturer’s protocol. RNA quality and concentration were assessed using a Qubit RNA HS Assay Kit (Thermo Fisher Scientific, USA) and Agilent 2100 Bioanalyzer. A total of 500 ng of high-quality RNA was used as input for Direct RNA Sequencing using the Oxford Nanopore SQK-RNA002 kit, following the manufacturer’s protocol. Briefly, sequencing adapters were ligated to the RNA molecules at the 3′ poly(A) tail, and reverse transcription was performed to stabilize RNA during sequencing. The prepared library was loaded onto an R9.4.1 flow cell (FLO-MIN106) and sequenced using the MinION Mk1B platform with MinKNOW software for data acquisition and live basecalling. Sequencing was conducted for 24 hours to obtain a sufficient number of long reads for transcriptome assembly.

### Transcriptome assembly and gene model prediction

For Nanopore RNA sequencing reads, we performed long read filtering and trimming using NanoFilt with parameter (-q 10 -l 500) (14) and de novo assembly using RNA-Bloom2 (15). Next, we performed gene model (protein-coding gene) prediction in each reference transcriptome using TransDecoder (https://github.com/TransDecoder/TransDecoder). Finally, we used BUSO pipeline (13) to assess the quality of reference transcriptome assemblies.

### Phylogenetic tree construction and divergence time estimation

We included the genomes and reference transcriptomes with more than 90% identified complete BUSCO genes. In addition, we downloaded the genome of striped bark scorpion *Centruroides vittatus* (GCF_030686945.1) for phylogenetic tree construction as the requirement of an outgroup species. BUSCO complete gene repertoire includes two categories, such as “Complete-Single-Copy (S)” and “Complete-Duplicated (D)”. We extracted the single-copy BUSCO genes “(S)” from 27 genomes and 121 reference transcriptomes, and identified one-to-one single-copy genes across 148 species using OrthoFinder (16). Further, we used a phylogenomic approach to reconstruct the phylogeny of 148 spider species and an outgroup scorpion species, based on a dataset of amino acid (AA) sequences corresponding to a pooled set of 1:1 single-copy genes. We performed AA sequence alignment using MAFFT (17), removed gaps using trimAL (18), and assembled a concatenated dataset that included all 1:1 single-copy genes with a minimum length of 200 AA. Finally, we used ModelFinder (19) to determine best-fit model of sequence evolution and constructed the maximum likelihood (ML) phylogenetic tree using IQ-TREE2 (20) with 1000 bootstrap replicates.

We retrievaled the documented divergence time between spiders on Time Tree database (https://timetree.org/). We estimated the divergence time for all nodes on the phylogeny using treePL (21). Finally, we used iTOL v6 (https://itol.embl.de/) to visualize the phylogenetic tree and divergence time.

### Orthologous group identification

Building on BUSCO gene repertoire which only included curated single-copy genes from OrthoDB database (https://www.orthodb.org), we employed FastOMA (22) pipeline to extend and further explore orthologous relationship of protein-coding genes across 148 spider species. For each 1:1 ortholog pair, the best-fit genes with longest coverage and highest similarity associated with newly generated orthologous groups (OGs) for each species were selected for further evolutionary analysis.

### Annotation of opsin, retinal determination and phototransduction pathway genes

We identified opsins, retinal determination and phototransduction pathway genes in genomes or representative high-quality reference transcriptomes of spider species with 0, 2, 4, 6, 8 eyes. Specifically, we searched for a previously compiled gene set including Arr, DAGK, Gprk1, Gprk2, rdgC, rdgB, PKC, PLC, Gα, Gβ, Gγ, trp, r-opsin, c-opsin, peropsin, rh1, rh2, rh3, rh4, ato, otd, Pax6, so/Six1, Six3, dac and eya (4, 23) in spiders. First, we linked the annotation of newly curated orthologs and the previously compiled gene set, and assigned orthologs to each spider visual system associated genes. Second, we used the protein sequences of compiled genes as queries to perform genome-wide or transcriptome-wide search using Blast-P or tBlast-N, and further annotated the candidate genes using NCBI-Blast with Non-redundant protein sequences database (NR).

### Hypothesized evolutionary transitions

By examining the spider tree of life (24–28) and instances of eye numbers at family or genus level, we hypothesized that evolutionary transitions from 8 eyes to 6, 4, 2, or 0 eyes may include multiple routes. Specifically, spider species in most of families have 8 eyes which was considered as ancestral trait. At family level, we hypothesized the evolutionary transition from 8 eyes to 2 eyes as route 1, including species in family Caponiidae; transition from 8 eyes to 4 eyes as route 2, including species in family Symphytognathidae; transition from 8 eyes to 6 eyes as route 3, including species in families Leptonetidae, Scytodidae, Sicariidae, Oonopidae and Leptonetidae. At genus level, we hypothesized the evolutionary transition in *Leptonetela* spider species of family Leptonetidae from 6 eyes to 0 eyes as route 4 (Fig. 2A).

### Analysis of nucleotide substitution rate

To test whether pattern of molecular evolution was associated with evolutionary transition of visual systems in spiders, we detected the changes in nucleotide substitution rate (dN/dS, ω) and selective strength (relaxation or intensification) acting on each hypothesized evolutionary route. Specifically, we prepared the codon alignments of shared orthologs by 148 spider species, which which derived from amino acid alignments and corresponding DNA sequences using PAL2NAL v.14 (-no gap) (29). We retained codon alignments with a minimum length of 50 codons, and prepared the corresponding tree for each ortholog by pruning the genome-scale phylogeny using R package, phytools (30). Finally, we used RELAX (31) to test for relaxation or intensification of selective pressure associated with each evolutionary route (Fig. 2A). Briefly, RELAX distinguishes between the signals by modeling how codons with different ω categories (ω□>□1 and ω□<□1) respond to a single selection intensity parameter K. Relaxation of selection would push all ω categories toward 1, while intensification of selection would pull all ω categories away from 1. K > 1 indicates the signature of intensified selection, whereas K < 1 indicates a relaxed selection strength at focal foreground branches (i.e., spider species with 2 eyes) (32, 33). We employed a log-likelihood ratio test (LRT) to compare the supports for the null model (K = 1) and the alternative model (K > 1 or K < 1), which further corrected P values using the Benjamini-Hochberg method to control for multiple comparisons. We defined the genes under parellel relaxation which showed K < and adjusted *P* < 0.05, while genes with K >1 and adjusted *P* < 0.05 are considered to be under parellel intensification.

### Analysis of positive selection

To determine whether positive selection was associated with evolutionary transition of visual systems in spiders, we detected the signal of positive selection in focal foreground branches (i.e., species with 2 eyes) relative to their ancestral branches of species with 8 eyes using Branch-site Unrestricted Statistical Test for Episodic Diversification for PHenotype (BUSTED-PH), which is a method to test for evidence of episodic diversifying selection associated with a specific phenotype/trait (34). Similarly, we used the codon alignments and gene trees for each shared ortholog as described above. We used LRT to calculate adjusted *P* values for focal foreground branches (adjusted *P* value-foreground), background branches (adjusted P value-background), and difference between focal foreground and background (adjusted *P* value-diff) following Benjamini–Hochberg adjustment. We defined genes with significant signals of parallel positive selection in invasive branches (adjusted *P*_foreground_ < 0.05) and difference between foreground and background branches (adjusted *P*_diff_ < 0.05), while no significant signals in nonsocial branches (adjusted *P*_background_ < 0.05) as positively selected genes in foreground spiders.

## Data availability

The raw sequencing reads have been deposited in NCBI under the project PRJNA1212500. All scripts required to perform all analyses are publicly available on GitHub (https://github.com/jiyideanjiao/Spider_Vision).

## Acknowledgments

This work was funded by the Special Investigation and Classification of Invertebrates from Yintiaoling Nature Reserve (CQS24C00333), the Science Foundation of School of Life Sciences SWU (20232008071901 and 20212020110501) to Z.S.Z and L.Y.W.

## Author contributions

C.T. and Z.S.Z designed research; C.T., Z.F., and L.Y.W. performed research; C.T. analyzed data; and C.T. wrote the paper.

